# Short-term intermittent hypoxia therapy promotes gliogenesis in a rat model of middle cerebral artery occlusion (MCAO) stroke

**DOI:** 10.1101/2023.06.26.546509

**Authors:** Syed Aasish Roshan, Dharani Gunaseelan, Swaminathan K Jayachandran, Mahesh Kandasamy, Muthuswamy Anusuyadevi

**Affiliations:** Molecular Neuro-Gerontology Laboratory, Department of Biochemistry, School of Life Sciences, Bharathidasan University, Tiruchirappalli-620024, Tamil Nadu, India; Drug Discovery and Molecular Cardiology Laboratory, Department of Bioinformatics, School of Life Sciences, Bharathidasan University, Tiruchirappalli-620024, Tamil Nadu, India; Laboratory of Stem Cells and Neuroregeneration, Department of Animal Science, School of Life Sciences, Bharathidasan University, Tiruchirappalli -620024, Tamil Nadu, India; University Grants Commission-Faculty Recharge Program (UGC-FRP), New Delhi 110002, India

**Keywords:** Neurogenesis, intermittent hypoxia, stroke, MCAo, DCX, GFAP

## Abstract

**Aim:** After focal cerebral ischemia, intermittent hypoxia therapy (IHT) could be used as a non-invasive method to stabilize and stimulate neurogenesis in the innate stem cell niche in the brain, and disrupt the glial scar around the infarct to increase neuroblast migration in the striatal infarct area.

**Methods:** We induced focal cerebral ischemia in Wistar albino rats using the MCAo model. A week later, animals were subjected to intermittent hypoxia (12%O_2_, 4hr/day) for a period of 14 days. Post-treatment analysis of functional recovery and cellular regeneration was done using immunofluorescence analysis of multiple neuronal cell markers including Doublecortin (DCX), Nestin, and Vimentin among others.

**Results:** Observations of GFAP-positive cells revealed that IH treatment facilitates gliogenesis in the infarct striatal region of a rat model of MCAo stroke. The percentage of DCX and GFAP double-positive cells was increased in the IH-treated group. Also, there was a significant difference in the morphology of vimentin-positive cells and microglia cells between the stroke groups.

**Conclusion:** These outcomes suggest that exposure of MCAo stroke-affected rats to intermittent hypoxia results in an increase in migrated neuroblasts resulting in a subsequent altered glial scar integrity in the infarct region, thus suggesting an alternative non-invasive method against the common stem cell transplant techniques, to increase endogenous neuroblasts in the infarct area after stroke.

## 1) Introduction

Ischemic stroke is caused by the blockade of an artery supplying the brain, predominantly the Middle Cerebral Artery (MCA). Loss of cerebral blood flow during an ischemic stroke causes a deprivation of oxygen and glucose in neurons(Campbell et al 2019). Consequently, a cascade of cellular and molecular events like excitotoxicity, oxidative stress, inflammation, necrosis, and apoptosis is triggered in the brain’s infarct zones, leading to neuronal cell death, and disruption of neural circuits (Kuriakose et al 2020). This severe loss of cellular mass, particularly neural, in the striatum, can cause severe impairments including paralysis, difficulty with speech or language, memory problems, and other cognitive disruptions (Chugh 2019). Hence targeting regeneration using neuroplasticity and neurogenesis is the most sought after for tissue regeneration in the damaged ischemic infarct. The neuropathogenesis of ischemic stroke has been shown to be highly associated with abnormal angiogenesis, impaired neurogenesis, defects in neuroplasticity, and an altered neurotrophic milieu in the brain. Especially, studies in rodents and humans have shown stroke to induce reactive neurogenesis in the SVZ and the migration of reactive neuroblasts toward the peri-infarct regions (Cuartero et al 2021, Marques et al 2019). From four days after the stroke to six weeks, increased neuroblast proliferation has been evident in the ipsilateral subventricular zone (SVZ) of the brain. Notably, migration of the newly generated neuroblasts towards the infarct region can be sustained for a limited period as the majority of the reactive neuroblasts appear to undergo apoptosis in the post-stroke brain. Moreover, a limited number of reactive neuroblasts manage to migrate, integrate, and survive, thus functional neurogenesis and neuroplasticity appear to be impaired in the brain of subjects with stroke (Lindvall 2015). As a result, spontaneous functional recovery has been not achieved due to the meager replacement of the lost neurons. One of the characteristic modulations of post-ischemic stroke is the prevalence of reactive astrogliosis and reactive microgliosis in the infarct hemisphere. The reactive astrocytes can form a dense cluster of cells, known as a glial scar, around the site of injury, which can prevent the migration of neural progenitor cells and the integration of new neurons into the existing neural circuitry(Pekny et al 2019). Glial scar is always considered to be one of the primary factors inhibiting neuroblast migration and survival (Wang et al 2018).

Intermittent hypoxia (IH) is an intervention where the animals are exposed to alternate periods of normoxia and hypoxia, mimicking the biomechanics of exercise (Brocherie 2020). This repeated cycle triggers various physiological responses, such as increased production of red blood cells, improved blood flow, and enhanced endurance, which are similar to the benefits seen with exercise training (Rybnikova 2022). IHT has shown that its increased neurogenesis has been attributed to the activation of molecular pathways that promote the proliferation of neural stem cells(Zhu 2010). Additionally, intermittent hypoxia has been shown to regulate the release of growth factors and cytokines that play a critical role in improvingneurogenesis(Khuu et al 2021). Furthermore, intermittent hypoxia has been shown to increase the expression of genes involved in neuroplasticity and synaptic formation, leading to improved cognitive function. IH has been shown to improve spatial learning and memory impairments in ischemia-induced stroke conditions through pMAPK and HIF-1α mediated hippocampal neurogenesis in association with the expression of c-Fos (Baillieul 2017). However, there is a huge lacuna where the fates of multiple cells in both SVZ and the infarct are completely unstudied under these conditions. Also, the effect of IH on glial scar modulation has never been reported before. Hence this study addresses how IHT-mediated modulations to neurogenesis actually contribute to neural proliferation and migration after ischemic stroke in an MCAo rat model.

## 2) Materials and Methods

### 2.1 Animals

Adult (16-week-old, 250g-330g) male Wistar albino rats were used in the study. Animals were procured from Liveon Bioloabs, Tumkur, and housed in Bharathidasan University animal housing facility. The 12-hour dark-light cycle was maintained, with ad libitum food and water availability. All experiments were conducted in accordance with the approval of the Institutional Animal Ethics Committee (IAEC) (Ref No: BDU/IAEC/P07/2019, November 11, 2019), under the regulation of the Committee for the Purpose of Control and Supervision of Experiments on Animals (CPCSEA), India. The rats were randomly divided into four groups: Sham group, Sham with IH treatment group, MCAo group, and MCAo with IH treatment group.

### 2.2 Animal model of Middle cerebral artery occlusion (MCAO) stroke model

The right middle cerebral artery (MCA) was occluded by the placement of a filament (Doccol Inc, USA) at its origin for a period of 90 minutes allowing reperfusion thereafter (Zhang et al., 1997). For sham operations, instant reperfusion was allowed.

### 2.3 TTC staining

This assay helps in determining the infarct region created by the stroke model. 24 hours after MCAo surgery, the animal was sacrificed, and the brain was removed and cut into 2-5mm slices using a rat brain matrix. The slices were treated with 2% TTC solution for 30 minutes. Later the tissues were fixed using PFA and images were taken.

### 2.4 Intermittent Hypoxia Treatment

Rats were allowed to recover for 7 days after surgery(Tsai et al,2008), after which animals were subjected to intermittent hypoxia (11%-14%O2) exposure using a hypoxic chamber for 4 hours per day for a period of 14 days.

### 2.5 Immunofluorescence

Rats were anesthetized, perfused, and later brain was isolated, and fixed in 4% paraformaldehyde. Post fixation, rat brains were processed in sucrose solutions and embedded in the Optimal Cutting Temperature (OCT) compound, before cryosectioning. 40µm thick free-floating sections were subjected to an immunostaining protocol. The following antibodies and their dilutions were used: Rabbit anti-NeuN (1:100, CST), Rabbit anti-Vimentin (1:250, CST), Rabbit anti-GFAP (1:250, CST), Rabbit anti-IBA1 (1:250, CST), Rabbit anti-DCX (1:250, CST), Mouse anti-DCX (1:250, Invitrogen), Mouse anti-Nestin (1:250, CST), Goat anti-rabbit Alexa 594(1:1000, CST), Goat anti-mouse Alexa 488(1:1000, CST). Sections were mounted with Invitrogen antifade mountant and observed using a fluorescent microscope.

### 2.6 Statistical Analyses

All data are expressed as the mean values ± SD. One-way ANOVAs also followed by Tukey’s post hoc test were used to assess the differences among groups. Student’s t-test was used when appropriate.

## 3) Results

### 3.1 Neuronal cell impairments after ischemia are selective and influence the post-ischemic recovery environment

Striatum is populated commonly by neurons, oligodendrocytes, astrocytes, and microglia among other cells. Following ischemia, the infarct zone encounters a severe loss of neuronal cells, evidenced through appropriate staining procedures. TTC staining of brain tissues collected 24 hours after the 90 min MCAo surgery revealed the infarct area localized to the ipsilateral striatum in MCAo groups (S.Fig-1). In this study, 21 days after ischemia, cresyl fast violet staining reveals severe loss of neurons (Coin-shaped) in MCAo animals(Fig-1-A-D). This evidence is also supported by NeuN staining to reveal severe loss of neurons in the infarct core of the ipsilateral striatum(Fig-1-E). Similar to the loss of neurons, loss of oligodendrocytes, and myelination are also evident in the infarct striatum, observed through immuno-staining against CNPase(Fig-1-I-L). These changes are observed throughout the MCAo groups.

**Figure-1:**
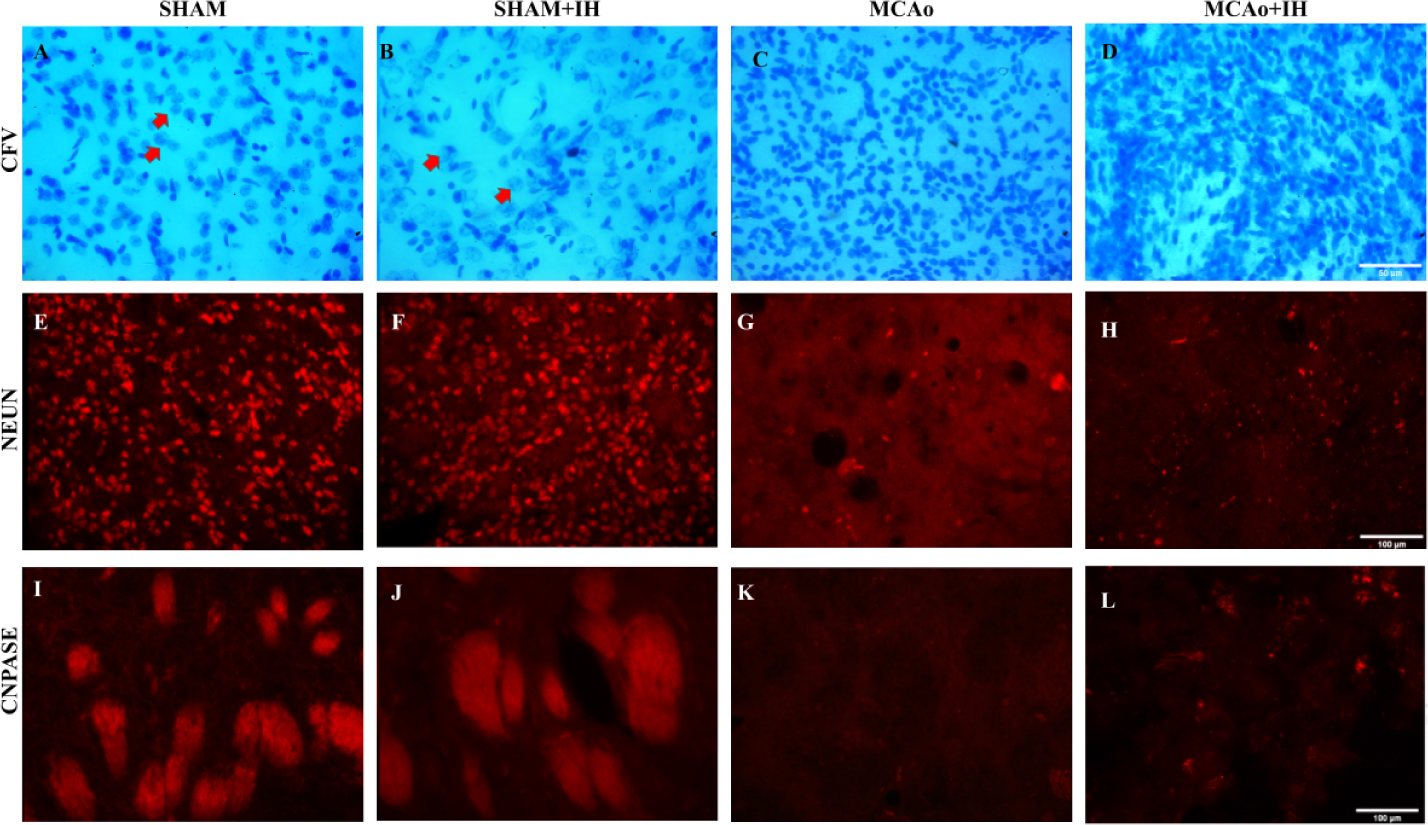
Neuronal cell impairments after ischemia are selective and influences post-ischemic recovery environment: A-D) CFV staining of tissue shows loss of neurons in the striatum of MCAo groups(C), and MI group(D). E-H) NeuN positive cells in the Stroke infarcted striatum of G) MCAo, and H) MI rats. I-L) CNPase positive cells in the Stroke infarcted striatum of K) MCAo, and L) MI rats.

### 3.2 Intermittent hypoxia alters post-ischemic reactive astrocytosis, as well as reactive microgliosis in the stroke infarct

Investigating the status of other neuronal cells, like astrocytes and microglia was carried out through immunostaining against GFAP, and IBA1. In the MCAo group, reactive astrocytosis is observed but restricted to the infarct boundary, and astrocytes were severely reduced in the infarct core. However, in the M+I group, reactive astrocytosis is observed in both the infarct core as well as the boundary(Fig-2-A-E). In aspects of microglia, IBA1 staining revealed a significant increase in reactive microgliosis throughout the infarct in both MCAo groups(Fig-2-F-J). However, the difference in microglia morphology was significantly evident, showing a higher number of amoeboid microglia, and a lower number of activated microglia in the MCAo group, whilst showing a higher number of activated microglia, and a lower number of amoeboid microglia in M+I group.

**Figure-2:**
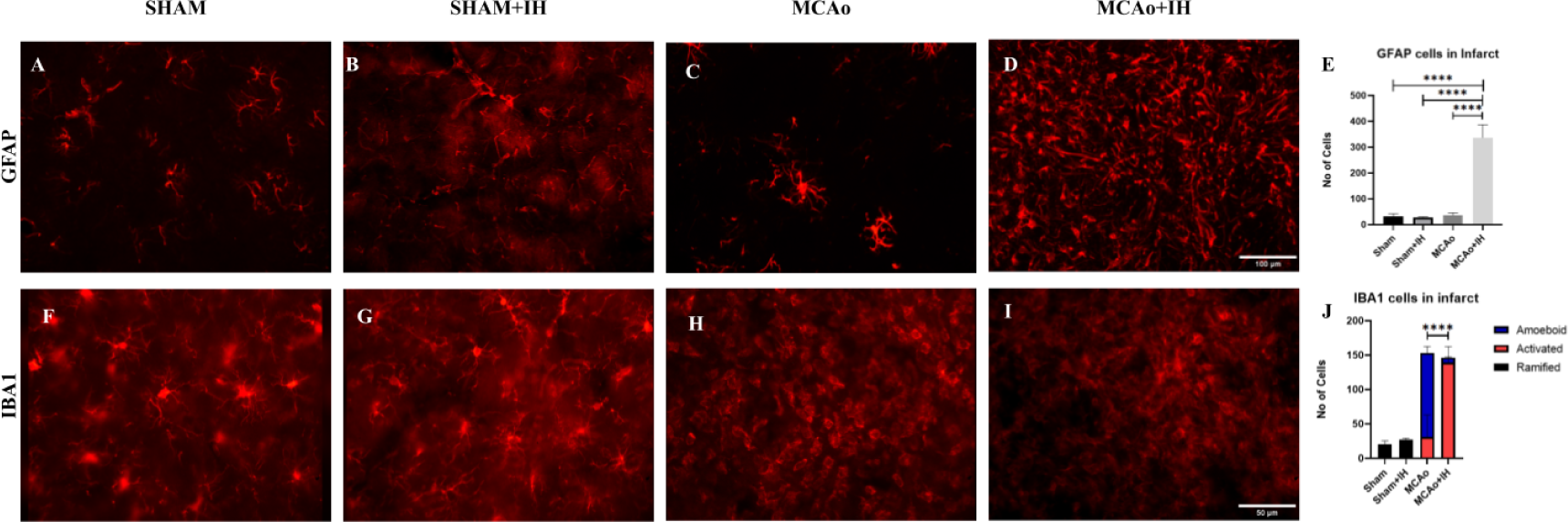
Intermittent hypoxia alters post-ischemic reactive astrocytosis, as well as reactive microgliosis in the stroke infarct. A-D) GFAP positive cells in the Stroke infarcted striatum of A) Sham, B) SI group, C) MCAo group, and D) MI rats. E) Quantitative analysis of GFAP positive Round Shaped Cells in the infarct striatum. There is a significant increase in GFAP-positive cells in the infarct core area of the MI group compared to MCAo.(One-way ANOVA, P-Value <0.0001). F-I) IBA1 positive cells in the Stroke infarcted striatum of A) Sham, B) SI group, C) MCAo group, and D) MI rats. J) Quantitative analysis of the three morphologies of IBA1 positive Cells in the infarct striatum. There is a significant increase in Activated IBA1 cells in the infarct core area of the MI group compared to MCAo.(One-way ANOVA, P-Value <0.0001).

### 3.3 Intermittent hypoxia alters ischemia-mediated stemcell proliferation in SVZ

To observe the characteristic ischemia-induced alterations in SVZ proliferation and migration, markers such as Ki67, Nestin, Vimentin, and Doublecortin(DCX) were assessed. Analysis of the proliferation marker Ki-67 revealed increased proliferation of stem cells in the MCAo groups in the SVZ. In comparison to the MI group, the MCAo group showed comparatively more proliferation(S.Fig-1). Evaluation of Nestin, a marker of neuroepithelial cells, revealed two distinct morphologies, where round-shaped cells were prominently dispersed throughout the walls of SVZ in all groups. However, process-shaped nestin-positive cells are present only in the MCAo groups(Fig-3-A-D). Assessment of vimentin, a marker of radial glia, also revealed multiple morphologies of inactive round-shaped cells and active elongated cells. However, active elongated vimentin cells are almost absent in sham groups, while their numbers are prominently increased in MCAo groups in comparison to the MI group(Fig-3-F-J). Analyzing DCX-positive cells in SVZ, revealed that when compared with the MCAo group, DCX-positive cells are slightly reduced in the MI group(Fig-3-K-O).

**Figure-3:**
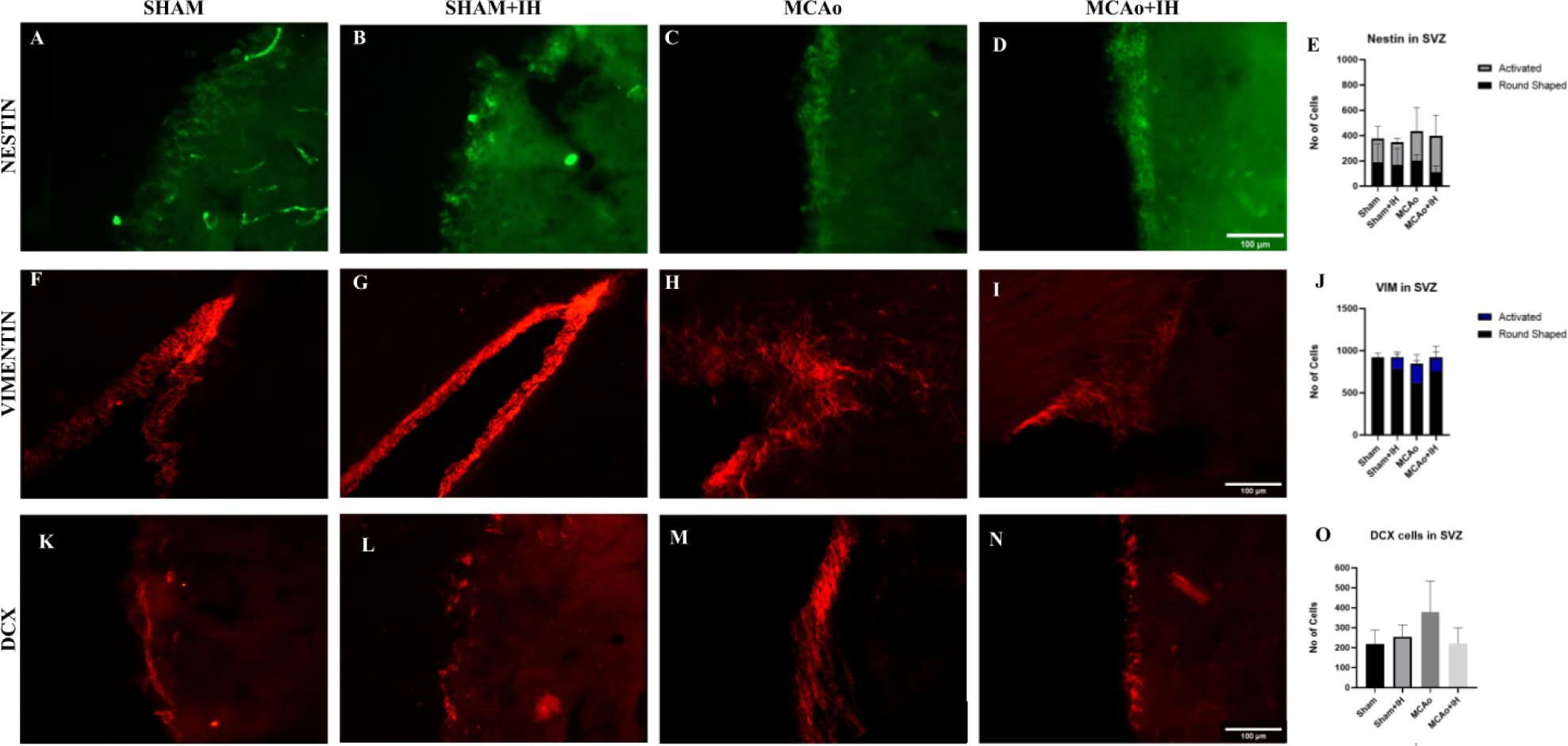
Intermittent hypoxia alters ischemia-mediated stemcell proliferation in SVZ: A-D) Nestin positive cells in the SVZ of A) Sham, B) SI group, C) MCAo group, and D) MI rats. E) Quantitative analysis of different morphologies of Nestin-positive Cells in the V-SVZ. F-I) Vimentin-positive cells in the SVZ of F) Sham, G) SI group, H) MCAo group, and I) MI rats. J) Quantitative analysis of different morphologies of VIM positive Cells in the SVZ. K-N) Doublecortin-positive cells in the SVZ of K) Sham, L) SI group, M) MCAo group, and N) MI rats. O) Quantitative analysis of different morphologies of DCX positive Cells in the SVZ.

### 3.4 Intermittent hypoxia increase Neuronal stem cell migration and survival in the ischemic infarct core

In order to observe the migration, survival, and morphology of neuronal stem cells, markers such as Ki67, Nestin, Vimentin, and Doublecortin(DCX) were assessed. Evaluation of the Ki67 marker revealed, significant upregulated levels of cell proliferation in the infarct core in MI group when compared with MCAo group(Fig-4-A-E). Assessment of Nestin positive cells in infarct core revealed IH treatment has significantly upregulated cell numbers in MI group in comparison with MCAo group(Fig-4-F-J). Similarly assessment of vimentin positive cells in the infarct revealed increase in number of these cells in MI group in comparison with MCAo group(Fig-4-K-O). Further analysis of vimentin positive cells revealed two distinct morphologies, where the round cellbody shaped vimentin cells are significantly increased after IH treatment in MCAo groups. Also Intermittent hypoxia has promoted migration and survival of DCX positive neuroblasts in the ischemic infarct core(Fig-4-P-T).

**Figure-4:**
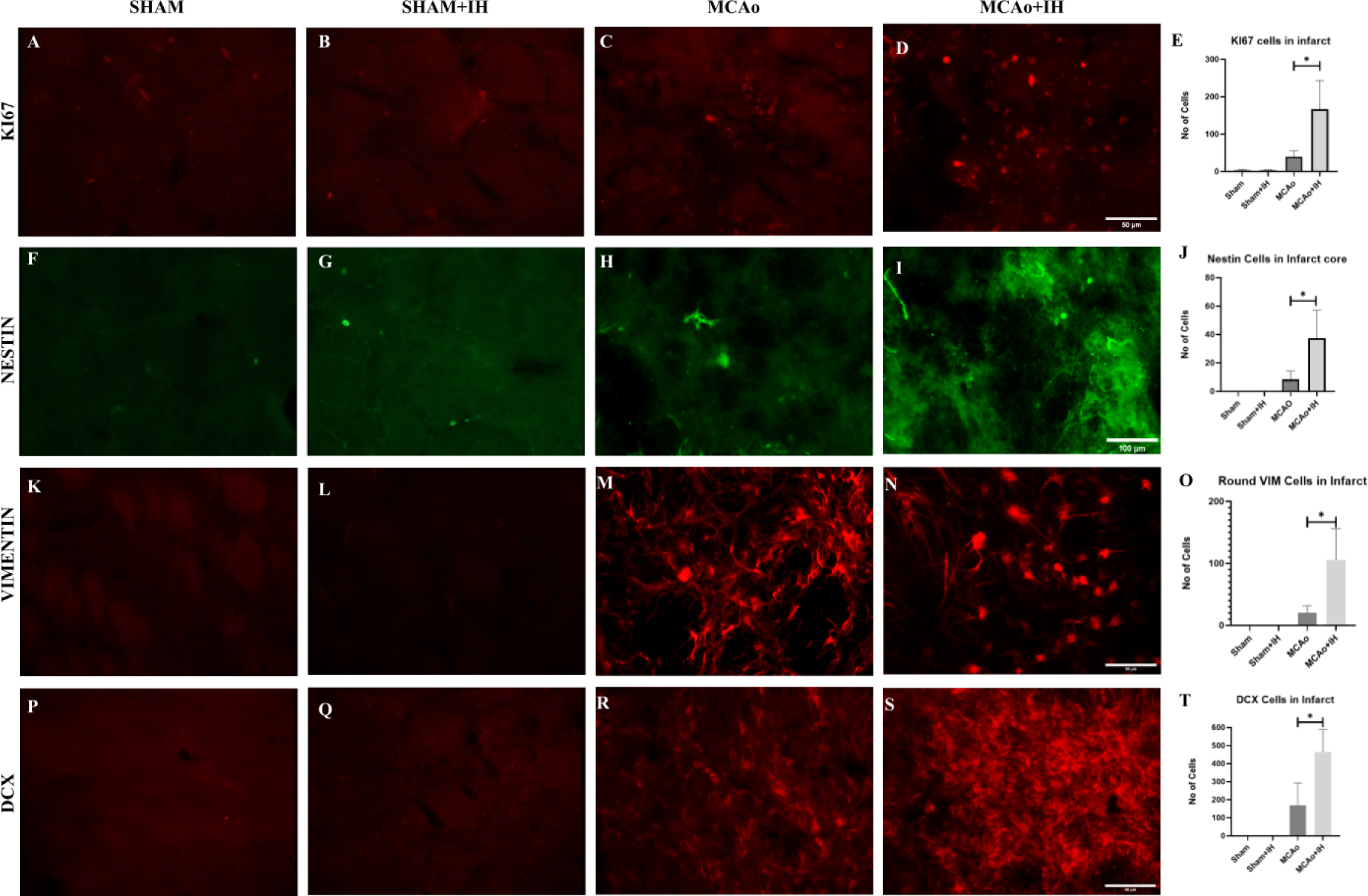
Intermittent hypoxia increase Neuronal stem cell migration and survival in the ischemic infarct core. A-D) KI-67 positive cells in the infarcted striatum of A) Sham, B) SI group, C) MCAo group, and D) MI rats. E) Quantitative analysis of KI-67-positive Cells in the infarcted striatum. There is a significant increase in KI-67 positive cells in the infarct core area of the MI group compared to MCAo.(Unpaired T test, P-Value= 0.0477). F-I) nestin-positive cells in the infarcted striatum of F) Sham, G) SI group, H) MCAo group, and I) MI rats. J) Quantitative analysis of Nestin positive Cells in the infarcted striatum. There is a significant increase in Nestin-positive cells in the infarct core area of the MI group compared to MCAo.(Unpaired T test, P-Value= 0.0369). K-N) Vimentin-positive cells in the infarcted striatum of K) Sham, L) SI group, M) MCAo group, and N) MI rats. O) Quantitative analysis of different morphologies of VIM positive Cells in the infarcted striatum. There is a significant increase in Migrated Round VIM-positive cells in the infarct core area of the MI group compared to MCAo.(Unpaired T test, P-Value= 0.0464). P-S) Doublecortin-positive cells in the infarcted striatum of K) Sham, L) SI group, M) MCAo group, and N) MI rats. T) Quantitative analysis of DCX positive Cells in the infarcted striatum. There is a significant increase in DCX-positive cells in the infarct core area of the MI group compared to MCAo.(Unpaired T test, P-Value= 0.0444).

### 3.5 Intermittent hypoxia disrupts glial scar characteristics in post-ischemic infarct

To understand variations in migration into the ischemic infarct, we assessed how glial scar is altered after stroke. GFAP-positive cells and nestin-positive cells were predominantly said to contribute to glial scar formation. Assessment of the GFAP-positive cells revealed that in the MCAo group, these cells are completely restricted to the infarct boundary and do not penetrate into the infarct core. However intermittent hypoxia treatment alters this restriction, enabling a continuous expansion of GFAP-positive astrocytes from the infarct boundary to the infarct core(Fig-5-A-F). Further assessment of Nestin positive cells reveals that intermittent hypoxia reduces the number of nestin positive cells in the infarct boundary, when compared to MCAo group, which has nestin positive cells lined throughout the infarct boundary(Fig-2-G-K).

**Figure-5:**
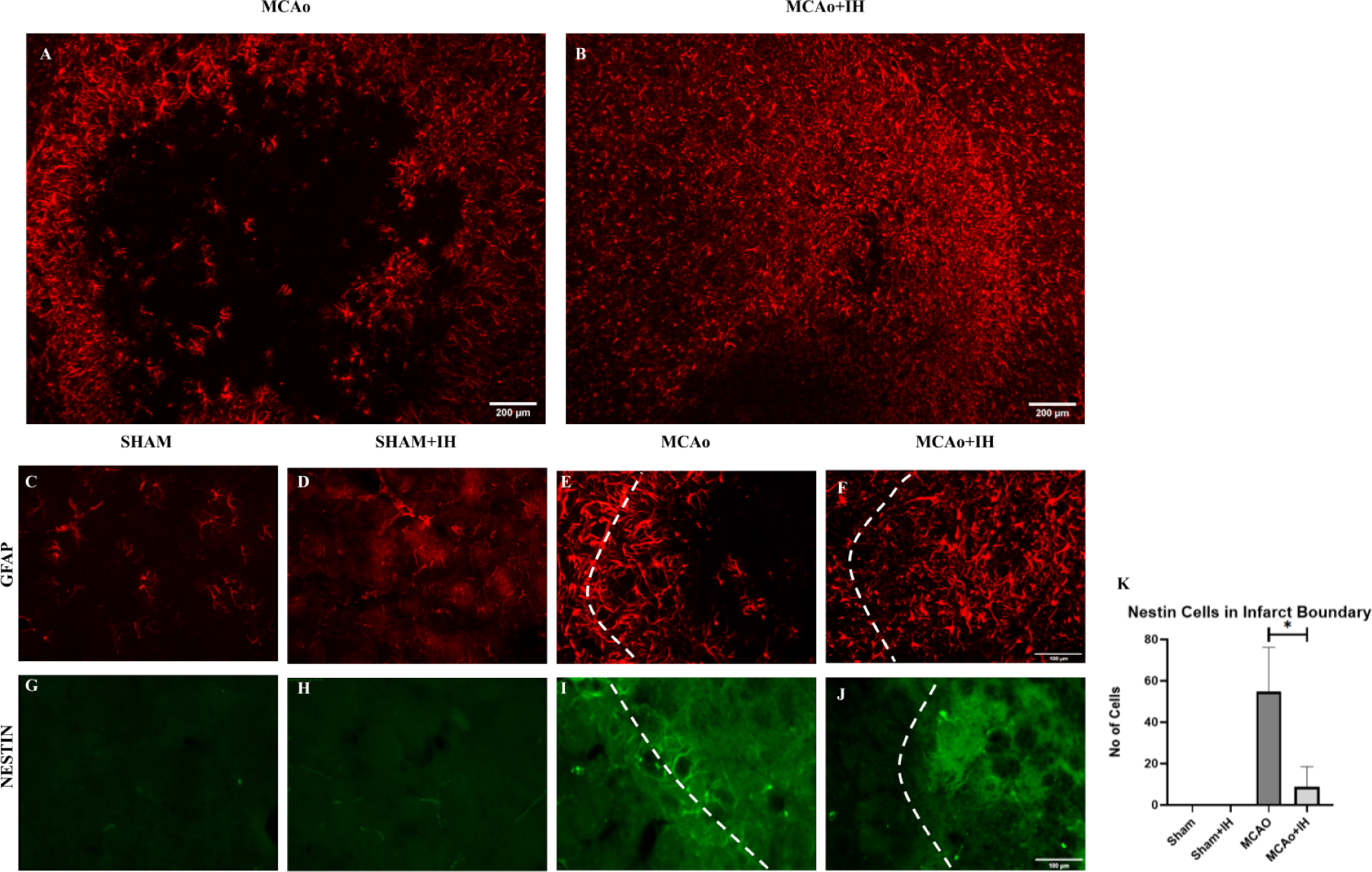
Intermittent hypoxia disrupts glial scar characteristics in post-ischemic infarct. A-F) GFAP positive cells in the Stroke infarcted striatum of A, E) MCAo group, B,F) MI rats, C) Sham, and D) SI group. G-J) Nestin positive cells in the boundary of the ischemic infarct of G) Sham, H) SI group, I) MCAo group, and J) MI rats. K) Quantitative analysis of Nestin-positive cells in the boundary of the ischemic infarct. There is a significant decrease in Nestin-positive cells in the boundary of the ischemic infarct of MI group compared to MCAo.(Unpaired t test, P-Value= 0.0280).

### 3.6 GFAP positive cells in the infarct near to the V-SVZ co-express DCX, but IBA1 positive cells donot co-express DCX

Since reactive astrogliosis and reactive microgliosis are the major sources of proliferation in the infarct zone, we analyzed if these cell types express DCX, and possibly contribute to neurogenesis. Colabeling study of GFAP/DCX revealed a significant overlay of GFAP-positive cells with DCX in the infarct boundary after stroke (Fig-6). Intermittent hypoxia increases the total number of GFAP/DCX co-labeled cells in the infarct zone. A similar assessment for IBA1/DCX revealed a meager confirmatory overlay of the cells (Fig-6).

**Figure-6:**
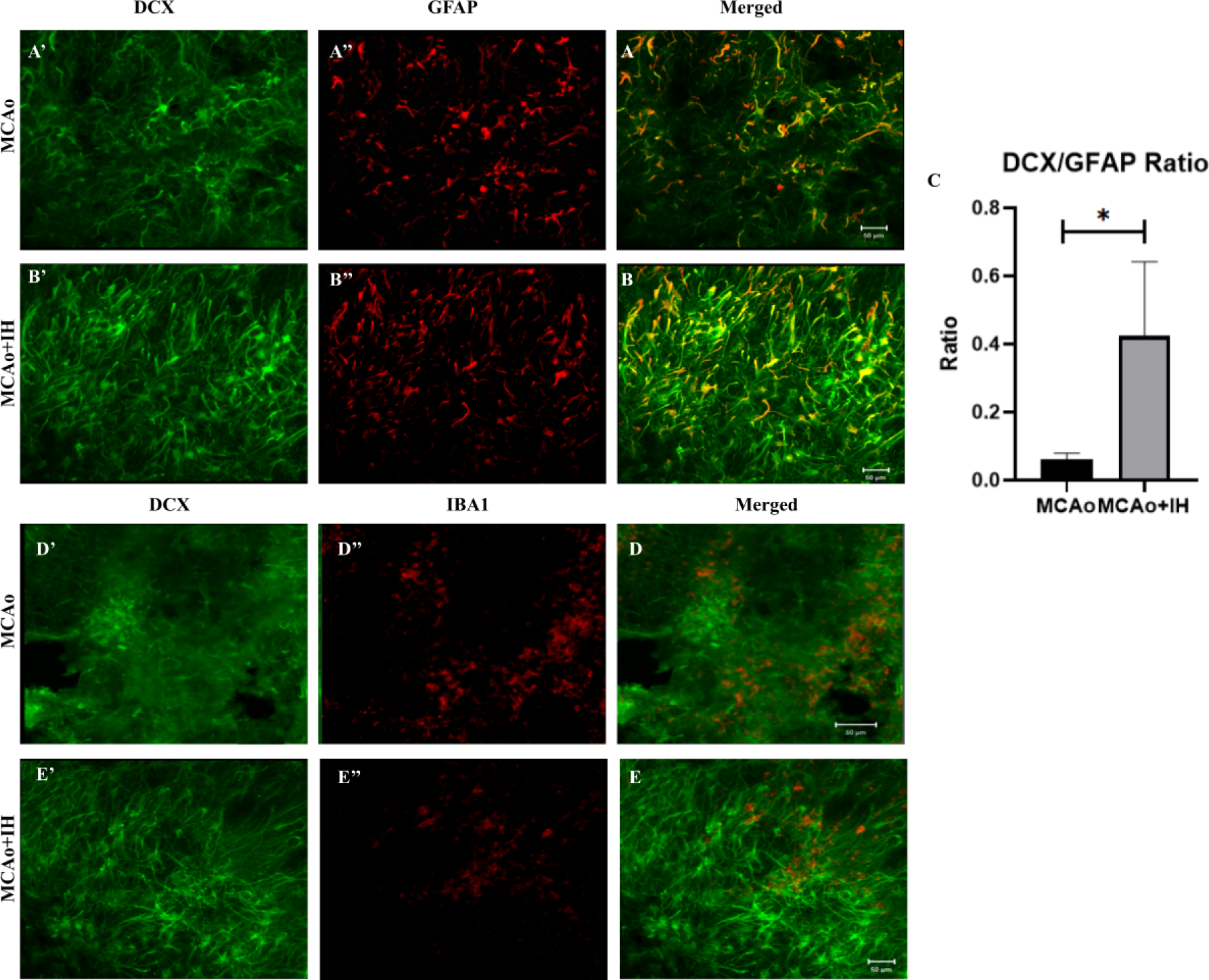
GFAP positive cells in the infarct near to the V-SVZ co-express DCX, but IBA1 positive cells donot co-express DCX. A) DCX positive cells (A’), and GFAP positive cells (A’’) in the boundary of the ischemic striatum of the MCAo group were merged(A). B) DCX positive cells (B’), and GFAP positive cells (B’’) in the boundary of the ischemic striatum of the MI group were merged(B). C) Quantitative analysis of the ratio between DCX, and GFAP double positive cells to total DCX positive cells was done. There is a significant increase in DCX-GFAP double-positive cells in the boundary of the ischemic infarct of the MI group compared to MCAo.(Unpaired t-test, P-Value= 0.0448). D) DCX positive cells (D’), and IBA1 positive cells (D’’) in the boundary of the ischemic striatum of the MCAo group were merged(D). E) DCX positive cells (E’), and GFAP positive cells (E’’) in the boundary of the ischemic striatum of the MI group were merged(E).

## 4) Discussion

The use of intermittent hypoxia therapy (IHT) for neuroprotection and neuroregeneration has long been a subject of interest in the field of neuroscience. However, the effects of IHT on the restoration of brain function after ischemic damage are not well understood. Previous studies have mainly focused on the neurogenesis of DCX-positive neuroblasts in the subventricular zone (SVZ) and factors that support them. Other studies have explored the effects of SVZ neurogenesis on the dentate gyrus (DG), leaving a gap in our understanding of neural stem cell situations in stroke infarct. Additionally, the modulation of glial barriers in post-stroke conditions following IHT remains largely unexplored. Therefore, in this study, we aimed to observe and address the alterations in the glial environment in post-stroke conditions following IHT, with the hope of contributing to a better understanding of the mechanisms underlying the neuroprotective and neuroregenerative effects of IHT.

Immediately after the stroke, the excitotoxicity-induced necrotic core, and surrounding apoptotic environment boundary region, decimates the local cellular environment in the ischemic infarct (Sekerdag et al 2018). This leads to a severe loss of neurons and their supporting cells in the striatal region. In our current study, observations at 21 days after stroke revealed that, till this time point, regeneration of neurons and oligodendrocytes were still absent in the ischemic core. Assessment of the same environment after IHT revealed no changes with neurons and oligodendrocytes through immunostaining against NeuN, and CNPase.

Reactive astrogliosis, the characteristic hallmark of post-stroke environment, has produced many GFAP astrocytes, still, their location is restricted to the infarct boundaries, leaving the infarct core, completely vague of GFAP-positive astrocytes (Zhang et al 2018). The specific loss of astrocyte at the core has been attributed to oncosis(Chu et al 2007, Liu et al 1999). The glial scar primarily comprises reactive astrocytes, which are activated by various cytokines, growth factors, and extracellular matrix molecules (Wang et al 2018). These activated astrocytes migrate to the site of injury and undergo morphological changes, including hypertrophy and upregulation of intermediate filament proteins such as glial fibrillary acidic protein (GFAP)(Lagos-Cabré R et al 2020, Karve et al 2016). This leads to the formation of a dense network of astrocytic processes around the damaged tissue, which serves as a physical barrier to prevent further injury and inflammation (Fitch et al 2008). In addition to astrocytes, microglia also play a role in glial scar formation. Microglia are activated by various inflammatory signals and can release cytokines and chemokines that recruit other immune cells to the site of injury (Dong et al 2021). Thus this desolated region of the ischemic infarct is predominantly occupied by microglia cells, with reactive microgliosis helping with the increased proliferation of the same. By modulating GFAP +Ve reactive astrocyte arrangement, and migration into the infarct area, IH supports more migration, increased neurogenesis, and should support revascularization of the infarct area, thereby could promote recovery after stroke. A similar assessment of the microglia in the infarct site reveal significant upregulation of microglia cells in both MCAo groups were observed. However further analysis of the morphology of the microglia reveal, in the MCAo group, the cells exhibited a predominant amoeboid shape, while the M+IHT group predominantly had activated cell types in the infarct zone. The functional roles of microglia in affiliation with its morphology has been reported previously, but whether it fits the description of the cells observed here is up for discussion.

This creates the need to assess how the altered gliogenesis encountered in the infarct hemisphere after the IHT has modulated neuronal stem cell proliferation and migration into the infarct. Observation of changes in the SVZ stem cell niche revealed some interesting findings. In post-stroke conditions, to facilitate the recovery of cells lost, SVZ stem cell niche increases neural proliferation(Arviddson et al). Similarly both MCAo and M+I group have exhibited an increase in proliferation in the SVZ, with latter at a lesser rate when analysed on day 21. The neural precursor cells (NPCs) in the SVZ can be identified based on their morphology and the expression of specific proteins. The most abundant are the type B cells characterized by small round cell body, and expressing nestin, vimentin as integral filament proteins(Bernal et al 2018). Assessment of the neuroepithelial cell marker, nestin, which revealed that, in the SVZ, these cells exhibit two different morphologies, a dormant round shape, and an active elongated cell. The elongated cells are predominantly present in VSVZ where their transition to the RMS begins, indicating active migration of these cell types. The increased number of elongated cells in the MCAo groups is an active indication that stroke induced migration of the nestin positive cells have increased. One other observation revealed, in all groups, Medial SVZ tends to have the majority of the round-shaped cells, whilst the lateral SVZ has the process-shaped nestin cells, thus exhibiting region-dependent cell states/phases. Vimentin is an intermediate filament protein, famously studied for its role in cellular migration, and also considered a marker for radial glia (Nakagawa et al 2004, Miranda-Negron et al 2022). In sham animals, throughout the SVZ, the cells are round-shaped, indicating an inactive resting state, where the vimentin proteins are organized around the nucleus, giving off a characteristic coin-shaped appearance. Approaching the RMS in VSVZ, vimentin-positive cells exhibit an elongated or bipolar morphology, embarking on their migration journey towards, the olfactory bulb in their sham groups, and diverting towards the infarct zone in MCAo groups. IH treatment in MCAo results in less process-shaped VIM cells in SVZ, indicating lesser activation of the type B cells. Type B cells divide slowly and give rise to rapidly dividing type C cells which express markers such as doublecortin (DCX), which is a microtubule-associated protein that is specifically expressed in migrating and differentiating neuroblasts(Lim et al 2016). These neuroblasts migrate along the rostral migratory stream (RMS) towards the olfactory bulb. After stroke, subventricular-zone-derived neuroblasts are recruited to the neurovascular niche in the peri-infarct region. Subsequent evaluation of doublecortin-positive neuroblasts reveals that they are localized only to the lateral SVZ throughout the groups. And the number of DCX-positive cells in VSVZ is slightly reduced in the MI group in comparison to MCAo. This 21-day study thus shows a trend on how by limiting the type B and type C cell proliferation, IH treatment could prevent early SVZ niche depletion and could prolong long-term neurogenesis after stroke.

Post their proliferation and activation at the SVZ, successful migration and integration needs to be assessed at the site of infarct. One prime factor that could be affecting the migration should be the glial barrier formed at the ischemic boundary (Nicaise et al 2022, Xu et al 2020, Yang et al 2020). IHT severely modulates the cells occupying the glial scar, thus enabling better migration into the infarct core. Assessment of nestin-positive cells projected that the migrated nestin-positive cells were restricted to the ischemic boundary in the MCAo group, but IHT enabled further migration into the ischemic core. This in conjunction with the expansion of GFAP positive astrocyte population in the ischemic infarct in the M+IHT group, clearly indicates the disturbances of the barrier and subsequently promotes migration of other cell types into the infarct.

The effect of IHT on gliogenesis and recovery is significantly well exhibited at the infarct site. IHT seem to improve the local cellular proliferation, as evident by increase in Ki67 cells in the infarct. This local proliferation may mostly contribute to local microgliosis or astrogliosis, even with the possibility of other cells. Though nestin-positive stem cells, and GFAP-positive cells are restricted at the infarct boundary, other stemcells that are DCX-positive, and vimentin-positive manage to enter the infarct even in the MCAo group. This shows that a subset of neural precursors that are not nestin and GFAP positive exist, that can go past the glial scar, to occupy ischemic core. Since IHT has shown signs of glial barrier disruption, it has promoted DCX-positive neuroblast migration through them, thus allowing significantly higher accumulation of DCX positive neuroblasts inside the ischemic core. The increased number of DCX-positive cells inside the infarct post-IHT is a positive indication of regeneration.

Similarly in aspects of vimentin, IHT has increased the number of vimentin cells in the ischemic core. In addition, IHT has caused presence of more round shaped VIM cells inside the infarct when compared to the presence of more elongated cell shapes observed in the MCAo group. It is characteristic that during differentiation, vimentin-positive stem cells can undergo changes in morphology as they adopt the characteristics of their target cell type (Mendez et al 2010). Here this could be an indication of the migrated cells trying to undergo differentiation before integrating into the environment. The observed round shape, could indicate the vimentin-positive precursor differentiating more into neurons than oligodendrocytes because of their large nucleus and bipolar nature. Together, all these modulations point towards a better regenerative environment being produced in post-IHT conditions.

The study also focused on investigating whether reactive astrogliosis and reactive microgliosis, the primary sources of proliferation in the infarct zone, express DCX and contribute to neurogenesis. The findings indicated a significant overlap of GFAP-positive cells with DCX in the infarct boundary after stroke, indicating that these cells may play a role in neurogenesis. Furthermore, the study revealed that intermittent hypoxia increases the total number of GFAP/DCX co-labeled cells in the infarct zone, suggesting that intermittent hypoxia may enhance neurogenesis.

However, the co-labeling study of IBA1/DCX revealed a meager confirmatory overlay of the cells. This suggests that while reactive microglia may contribute to proliferation in the infarct zone, they may not play a significant role in neurogenesis. These findings suggest that reactive astrogliosis is a more important factor in promoting neurogenesis after stroke.

The study has important implications for the development of therapies to enhance neurogenesis after stroke. Targeting reactive astrogliosis may be a promising strategy for promoting neurogenesis and improving functional recovery after stroke. Furthermore, the findings highlight the potential of intermittent hypoxia as a therapeutic approach to enhance neurogenesis in the infarct zone. However, further research is needed to fully understand the mechanisms underlying these effects and to optimize therapeutic strategies.

## 5. Conclusion

In conclusion, the results of our study demonstrate the potential of intermittent hypoxia therapy (IHT) as a non-invasive method to promote cellular regeneration and functional recovery following focal cerebral ischemia. Our findings suggest that IHT can modulate the glial scar formation and stimulate neurogenesis in the innate stem cell niche of the brain, leading to increased neuroblast migration in the striatal infarct area. These observations imply that IHT could be a promising alternative to traditional stem cell transplant techniques to promote endogenous neuroblasts in the infarct area after stroke.

Moreover, our study highlights the potential translational value of IHT, as it is a relatively easy and low-cost method that can be readily implemented in clinical settings. The use of immunofluorescence analysis of multiple neuronal cell markers including DCX, Nestin, and Vimentin provided valuable insights into the cellular mechanisms underlying IHT-induced regeneration. Future studies could further investigate the long-term effects of IHT on functional outcomes and assess the safety and feasibility of this therapy in human patients.

Overall, our findings suggest that IHT could be a promising therapeutic strategy for stroke patients, and further research is warranted to fully understand its potential benefits and limitations.

## Contributions

Conceptualization: SAR, AM; data acquisition: SAR; analysis and interpretation of data: SAR, GE, DG, SKJ, MK, AM; writing–original draft preparation: SAR; writing–review and editing: SAR, SKJ, MK, AM; critical revision of the manuscript for intellectual content: SKJ, MK, AM; supervision: AM

## Acknowledgments

The authors would like to thank Mercy Priyadharshini for proofreading the manuscript and for helpful suggestions.

## Data Availability

The data supporting the findings of this study are available within the article and its supplementary material.

