## Supplementary material for "Short-term intermittent hypoxia therapy promotes gliogenesis in a rat model of middle cerebral artery occlusion (MCAO) stroke": S.Fig1

**Supplementary Figure-1:**

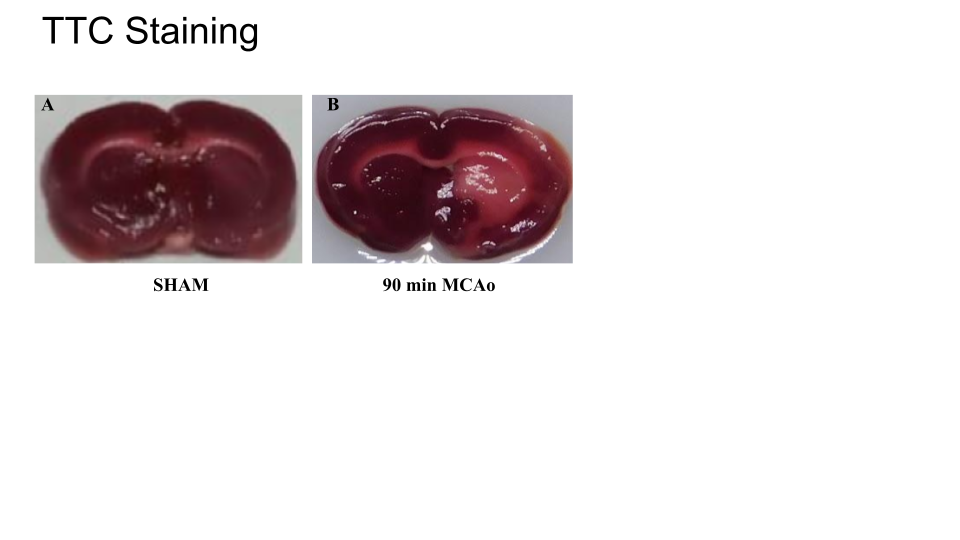

*S.Fig-1: TTC staining of the striatal region of rat brain compared among the sham(A), 90min MCAo occlusion(B).*
